# Microstructural Integrity of White Matter Moderates an Association Between Childhood Adversity and Adult Trait Anger

**DOI:** 10.1101/307637

**Authors:** M. Justin Kim, Maxwell L. Elliott, Tracy C. d’Arbeloff, Annchen R. Knodt, Spenser R. Radtke, Bartholomew D. Brigidi, Ahmad R. Hariri

**Author notes:** Correspondence: Justin Kim, PhD, Laboratory of NeuroGenetics, Department of Psychology and Neuroscience, Duke University, Durham, NC 27708, USA.

## Abstract

Amongst a number of negative life sequelae associated with childhood adversity is the later expression of a higher dispositional tendency to experience anger and frustration to a wide range of situations (i.e., trait anger). We recently reported that an association between childhood adversity and trait anger is moderated by individual differences in both threat-related amygdala activity and executive control-related dorsolateral prefrontal cortex (dlPFC) activity, wherein individuals with relatively low amygdala *and* high dlPFC activity do not express higher trait anger even when having experienced childhood adversity. Here, we examine possible structural correlates of this functional dynamic using diffusion magnetic resonance imaging data from 647 young adult men and women volunteers. Specifically, we tested whether the degree of white matter microstructural integrity as indexed by fractional anisotropy modulated the association between childhood adversity and trait anger. Our analyses revealed that higher microstructural integrity of multiple pathways was associated with an attenuated link between childhood adversity and adult trait anger. Amongst these pathways was the uncinate fasciculus, which not only provides a major anatomical link between the amygdala and prefrontal cortex but also is associated with individual differences in regulating negative emotion through top-down cognitive reappraisal. These findings suggest that higher microstructural integrity of distributed white matter pathways including but not limited to the uncinate fasciculus may represent an anatomical foundation serving to buffer against the expression of childhood adversity as later trait anger, which is itself associated with multiple negative health outcomes.

## Introduction

Experiencing adversity during childhood has multiple consequences on brain development, particularly that associated with the experience of negative affect (Tottenham, 2014). Functional and structural neuroimaging studies have highlighted corticolimbic brain regions associated with childhood adversity including exaggerated amygdala activity to threat-related facial expressions (Dannlowski et al., 2012), decreased intrinsic functional connectivity between the amygdala and ventromedial prefrontal cortex (Burghy et al., 2012), decreased orbitofrontal cortex volume (Hanson et al., 2010), and increased amygdala volume accompanied by elevated anxious temperament (Kuhn et al., 2016). This research to date has largely focused on the neural correlates of childhood adversity associated with the later experience of depression and anxiety (Nusslock and Miller, 2016; Casey et al., 2011).

In contrast, considerably fewer studies have examined brain regions that may link childhood adversity with the later expression of trait anger, which reflects an individual’s dispositional propensity for having a low threshold for feeling anger and perceiving a wide range of situations as frustrating (Spielberger, 1991). As such, trait anger has broad societal impact including increased risk for physical and mental illness, such as coronary heart disease and reactive aggression/violence, respectively (Chida and Steptoe, 2009; Bettencourt et al., 2006). Neuroimaging research has suggested that dysfunction in the corticolimbic circuit also may underpin trait anger and aggression (Davidson et al., 2000). For example, reduced functional connectivity between the amygdala and the prefrontal cortex has been mapped onto aggressive behavioral traits (Passamonti et al., 2008; Coccaro et al., 2007, but see also Beyer et al., 2014). Furthermore, a task-based activation study by our group has reported a positive correlation between trait anger and amygdala activity to angry facial expressions in men who are also high in trait anxiety, a pattern consistent with reactive aggression (Carré et al., 2012).

More recently, we reported that threat-related amygdala activity and executive control-related dorsolateral prefrontal cortex (dlPFC) activity jointly moderate a link between childhood adversity and trait anger in young adults, such that this association was attenuated for individuals with the combination of relatively low amygdala *and* high dlPFC activity (Kim et al., in press). Here, we aimed to expand upon these findings by utilizing diffusion magnetic resonance imaging (dMRI), which enables the quantification of the microstructural integrity of white matter fiber tracts (Basser and Pierpaoli, 1996). Based on the findings from the aforementioned research, we hypothesized that higher microstructural integrity of the uncinate fasciculus (UF), a major white matter fiber pathway connecting the prefrontal cortex with limbic areas including the amygdala (Ebeling and von Cramon, 1992), would buffer against the expression of higher trait anger associated with the experience of childhood adversity. Our hypothesis further reflects earlier work finding that higher microstructural integrity of the UF is associated with more frequent use of cognitive reappraisal to regulate negative emotions (Eden et al., 2015; Zuurbier et al., 2013), which reflects the top-down inhibition of amygdala activity by prefrontal circuits (Buhle et al., 2014). As a test of specificity, we performed exploratory analyses on other major white matter pathways across the whole brain.

## Materials & Methods

### Participants

A total of 647 undergraduate students (385 women, age range 18-22 years, mean age = 19.6 years) who successfully completed the Duke Neurogenetics Study (DNS) between January 25^th^, 2010 and November 12^th^, 2013 had available dMRI (two scans) and self-reported questionnaire data for our analyses. These participants were free of past or current diagnosis of a DSM-IV Axis I or select Axis II (borderline and antisocial personality) disorder assessed with the electronic Mini International Neuropsychiatric Interview (Lecrubier et al., 1997) and Structured Clinical Interview for the DSM-IV subtests (First et al., 1996), respectively. The DNS aims to assess the associations among a wide range of behavioral, neural, and genetic variables in a large sample of young adults. For the present study, we specifically focused on trait anger and its association with childhood adversity.

Prior to data collection, informed consent in accordance with the Duke University Medical Center Institutional Review Board was obtained from all participants. To be eligible for the DNS, all participants were free of the following conditions: 1) medical diagnoses of cancer, stroke, head injury with loss of consciousness, untreated migraine headaches, diabetes requiring insulin treatment, chronic kidney, or liver disease; 2) use of psychotropic, glucocorticoid, or hypolipidemic medication; and 3) conditions affecting cerebral blood flow and metabolism (e.g., hypertension).

### Self-Report Questionnaires

The Childhood Trauma Questionnaire (CTQ) was used to assess exposure to childhood adversity in five categories: emotional abuse, physical abuse, sexual abuse, emotional neglect, and physical neglect (Bernstein et al., 1997). The State-Trait Anger Expression Inventory (STAXI) was used to index trait anger (Spielberger, 1991). In addition to total scores, the two subscales of STAXI – angry temperament or the propensity to experience anger without being provoked and angry reaction or the propensity to experience anger in response to negative events - were also calculated. Trait anxiety was measured using the State-Trait Anxiety Inventory-Trait Version (STAIT), in order to test whether or not the present findings were specific to trait anger (Spielberger et al., 1988).

### Image Acquisition

Diffusion-weighted and high-resolution anatomical T1-weighted magnetic resonance imaging scans were acquired using an 8-channel head coil for parallel imaging on one of two identical research-dedicated GE MR750 3T scanners (GE Healthcare) at the Duke-UNC Brain Imaging and Analysis Center. Following an ASSET calibration scan, diffusion-weighted images were acquired across two consecutive 2-min 50-s providing full brain coverage with 2 mm isotropic resolution and 15 diffusion-weighted directions (echo time (TE) = 84.9 ms, repetition time (TR) = 10,000 ms, *b* value = 1,000 s/mm^2^, field of view (FOV) = 240 mm, flip angle = 90°, matrix = 128 × 128, slice thickness = 2 mm). High-resolution anatomical T1-weighted MRI data were obtained using a 3D Ax FSPGR BRAVO sequence (TE = 3.22 ms, TR = 8.148 ms, FOV = 240 mm, flip angle = 12°, 162 sagittal slices, matrix =256 × 256, slice thickness = 1 mm with no gap).

### Diffusion MRI Analysis

All dMRI data were preprocessed in accordance with the protocol developed by the Enhancing Neuro Imaging Genetics through Meta-Analysis consortium (ENIGMA; http://enigma.ini.usc.edu/protocols/dti-protocols/). Raw diffusion-weighted images were corrected for eddy current and aligned to the non-diffusion-weighted (b0) image using linear registration in order to correct for head motion. Volume-by-volume head motion was quantified by calculating the root mean square (RMS) deviation of the six motion parameters (three translation and three rotation components), for each pair of consecutive diffusion-weighted brain volumes. The resulting volume-by-volume RMS deviation values were averaged across all images, yielding a summary statistic of head motion for each participant, which were used as a covariate of no interest in all subsequent group level analyses. Next, following skull stripping, diffusion tensor models were fit at each voxel using the Diffusion Toolbox in FSL (Smith et al., 2004; Behrens et al., 2003), generating a whole-brain fractional anisotropy (FA) image for each participant. For each participant, an average of the FA images from each of the two scans were produced and these images were then subjected to a tract-based spatial statistics (TBSS) in FSL (Smith et al., 2006). TBSS analysis entails the realignment of each individual FA image to a standard FA template in Montreal Neurological Institute (MNI) space using nonlinear registration (FNIRT). Next, a group-mean FA image was generated and subsequently thinned at a threshold of FA > 0.2 to extract the mean FA skeleton, which is assumed to represent the center of the white matter fiber tracts common to the group. Then, individual FA images were projected onto the mean FA skeleton, searching for maximal FA values perpendicular to the skeleton. The resulting individual FA skeletons were used to extract average FA values for our *a priori* pathways of interest.

### Pathways-of-Interest Analysis

*A priori* pathways of interest (POIs) were taken from the Johns Hopkins University DTI-based white matter atlas, adhering to the ENIGMA protocol (Wakana et al., 2007). These POIs were used to mask each participant’s FA skeleton maps, and then the average FA values were extracted on a subject-by-subject basis. As the UF included in this atlas only represents a very small intermediary segment of the pathway, we used an UF POI with better coverage of the entire tract using the Johns Hopkins University White Matter Tractography Atlas (Mori et al., 2005). The left and right UF POIs were binarized in order to extract mean FA values from the left UF and right UF for each participant. Since we did not have *a priori* predictions regarding inter-hemispheric differences and to reduce the number of statistical tests, the extracted FA values were averaged across left and right hemispheres for further statistical analyses (see Table 1 for a full list of the 24 POIs).

**Table 1.** Moderating effects of white matter pathway strength on the association between childhood adversity on trait anger, controlling for the effects of age, sex, and head motion. **Abbreviations**: ACR (anterior corona radiata), ALIC (anterior limb of the internal capsule), BCC (body of the corpus callosum), CGC (cingulum-cingulate gyrus), CGH (cingulum-hippocampus), CP (cerebral peduncle), CST (corticospinal tract), EC (external capsule), FX (fornix), FXST (fornix-stria terminalis), GCC (genu of the corpus callosum), ICP (inferior cerebellar peduncle), ML (medial lemniscus), PCR (posterior corona radiata), PLIC (posterior limb of the internal capsule), PTR (posterior thalamic radiation), RLIC (retrolenticular part of the internal capsule), SCC (splenium of the corpus callosum), SCP (superior cerebellar peduncle), SCR (superior corona radiata), SFO (superior fronto-occipital fasciculus), SLF (superior longitudinal fasciculus), SS (sagittal stratum), UF (uncinate fasciculus).

| Tract | *b* (SE) | 95% CI | $\Delta R^2$ | $F_{(1,640)}$ | *p* |
| --- | --- | --- | --- | --- | --- |
| ACR | -2.57 (0.77) | [-4.09, -1.05] | 0.017 | 11.04 | 0.0009* |
| UF | -3.10 (1.02) | [-5.10, -1.10] | 0.014 | 9.25 | 0.0024* |
| SLF | -2.81 (0.95) | [-4.68, -0.94] | 0.013 | 8.68 | 0.0033* |
| GCC | -2.45 (0.88) | [-4.20, -0.76] | 0.012 | 8.00 | 0.0048* |
| EC | -3.14 (1.11) | [-5.33, -0.96] | 0.012 | 8.00 | 0.0048* |
| SFO | -1.83 (0.66) | [-3.12, -0.53] | 0.012 | 7.71 | 0.0056* |
| SCC | -2.57 (0.95) | [-4.43, -0.72] | 0.011 | 7.40 | 0.0067* |
| SS | -1.99 (0.75) | [-3.46, -0.52] | 0.011 | 7.06 | 0.0081* |
| PTR | -1.52 (0.65) | [-2.80, -0.24] | 0.008 | 5.41 | 0.0204<sup>§</sup> |
| ALIC | -2.09 (0.93) | [-3.92, -0.26] | 0.008 | 5.02 | 0.0254 |
| RLIC | -1.56 (0.78) | [-3.09, -0.03] | 0.006 | 3.98 | 0.0464 |
| PCR | -1.72 (0.88) | [-3.45, 0.01] | 0.006 | 3.82 | 0.0511 |
| SCR | -2.13 (1.11) | [-4.30, 0.04] | 0.006 | 3.73 | 0.0539 |
| BCC | -1.69 (0.89) | [-3.43, 0.05] | 0.006 | 3.64 | 0.0567 |
| CP | 1.42 (1.28) | [-1.10, 3.95] | 0.002 | 1.23 | 0.2679 |
| CGC | -0.63 (0.61) | [-1.83, 0.57] | 0.002 | 1.06 | 0.3040 |
| ML | -0.54 (0.62) | [-1.75, 0.68] | 0.001 | 0.75 | 0.3878 |
| CGH | -0.44 (0.56) | [-1.54, 0.67] | < 0.001 | 0.60 | 0.4383 |
| FXST | 0.50 (0.78) | [-1.02, 2.02] | < 0.001 | 0.42 | 0.5189 |
| FX | 0.23 (0.49) | [-0.72, 1.19] | < 0.001 | 0.23 | 0.6318 |
| CST | 0.29 (0.71) | [-1.10, 1.68] | < 0.001 | 0.17 | 0.6811 |
| PLIC | -0.45 (1.14) | [-2.67, 1.78] | < 0.001 | 0.15 | 0.6952 |
| ICP | -0.23 (0.72) | [-1.65, 1.18] | < 0.001 | 0.10 | 0.7467 |
| SCP | 0.05 (0.70) | [-1.32, 1.42] | < 0.001 | 0.01 | 0.9423 |

\* $q < 0.05$ , <sup>§</sup> $q < 0.05$ when whole brain FA was included as a covariate

\**q* < 0.05, ^§^*q* < 0.05 when whole brain FA was included as a covariate

#### Study Design and Statistical Analysis

PROCESS for SPSS (Hayes, 2013) was utilized within SPSS 21 (IBM Corp., Armonk, NY, USA) to test whether FA values of white matter pathways moderated the association between CTQ and STAXI scores (independent and dependent variables, respectively), while including age, sex, and head motion as covariates. We also report the changes in the main findings when global FA (i.e., average FA across all white matter tracts) was also included as a covariate. As per our *a priori* hypothesis, the initial moderation analysis focused on the UF; subsequently, the remaining 23 POIs were analyzed using identical procedures. Since 24 major fiber pathways were tested separately (i.e., 24 independent moderation models), a false discovery rate correction was imposed on the significance threshold (*q* < 0.05) to correct for multiple statistical tests (Benjamini and Hochberg, 1995). Finally, to assess the specificity of the present findings to trait anger, an additional set of *post hoc* analyses were performed with trait anxiety (STAI-T scores) as the dependent variable in the moderation models.

## Results

### Self-Report Questionnaire Results

Means and standard deviations for the self-report measures were as follows: STAXI (15.77 ± 4.19), CTQ (33.06 ± 7.88), STAI-T (37.07 ± 8.6). Scores for the STAXI subscales were as follows: angry temperament (5.24 ± 1.81) and angry reaction (7.87 ± 2.47). Scores for the CTQ subscales were as follows: emotional abuse (7.01 ± 2.54), physical abuse (5.98 ± 1.82), sexual abuse (5.24 ± 1.44), emotional neglect (8.34 ± 3.47), and physical neglect (6.48 ± 2.19). As expected, STAXI total scores were positively correlated with CTQ total scores (*r* = 0.15, *p* < 0.001).

### dMRI Results

Five out of 24 pathways examined showed significant negative correlations between FA and CTQ at *p* < 0.05, after controlling for age, sex, and head motion. These included the UF (*r* = −0.1, *p* = 0.011) as well as the superior cerebellar peduncle (*r* = −0.15, *p* = 0.0002); medial lemniscus (*r* = −0.14, *p* = 0.0006); inferior cerebellar peduncle (*r* = −0.09, *p* = 0.018); and anterior corona radiata (*r* = −0.08, *p* = 0.041). However, only association for the superior cerebellar peduncle and medial lemniscus (*q*s < 0.05) remained significant after correction for multiple comparisons. Three out of 24 pathways showed significant correlations with STAXI at *p* < 0.05 (posterior thalamic radiation: *r* = −0.12, *p* = 0.002; sagittal striatum: *r* = −0.1, *p* = 0.016; splenium of the corpus callosum: *r* = −0.08, *p* = 0.034), but none survived correction for multiple comparisons (both *q*s > 0.05).

### Moderation Analysis Results

The overall model including childhood adversity and microstructural integrity of the UF was significant in predicting trait anger (*R*^2^ = 0.04, *F*(6, 640) = 4.4, *p* = 0.0002). A significant interaction effect indicated that the FA of the UF moderated the association between CTQ and STAXI (*b* = −3.1, 95% confidence interval = [−5.1, −1.1], Δ*R*^2^ = 0.01, *p* = 0.0024). Follow up simple slopes analysis showed that the interaction was primarily driven by individuals with relatively higher FA of the UF, for whom the association between CTQ and STAXI total scores was attenuated (*b* = 0.01, 95% CI = [−0.52, 0.72], *p* = 0.75). The Johnson-Neyman calculation indicated that childhood adversity was significantly associated with trait anger only when UF FA was less than 0.41 standard deviations above the mean. However, a similar pattern was also found in 7 other pathways even when applying a false discovery rate-corrected threshold for multiple comparisons (*q* < 0.05; Figure 1, Table 1). These interactions were robust to the inclusion of global FA as a covariate in the model (i.e., the same 8 pathways remained significant, along with the posterior thalamic radiation). Finally, when the moderation models were tested with trait anxiety as the dependent variable, none of the 24 POIs showed a significant effect (all *q*s > 0.05).

**Figure 1.**
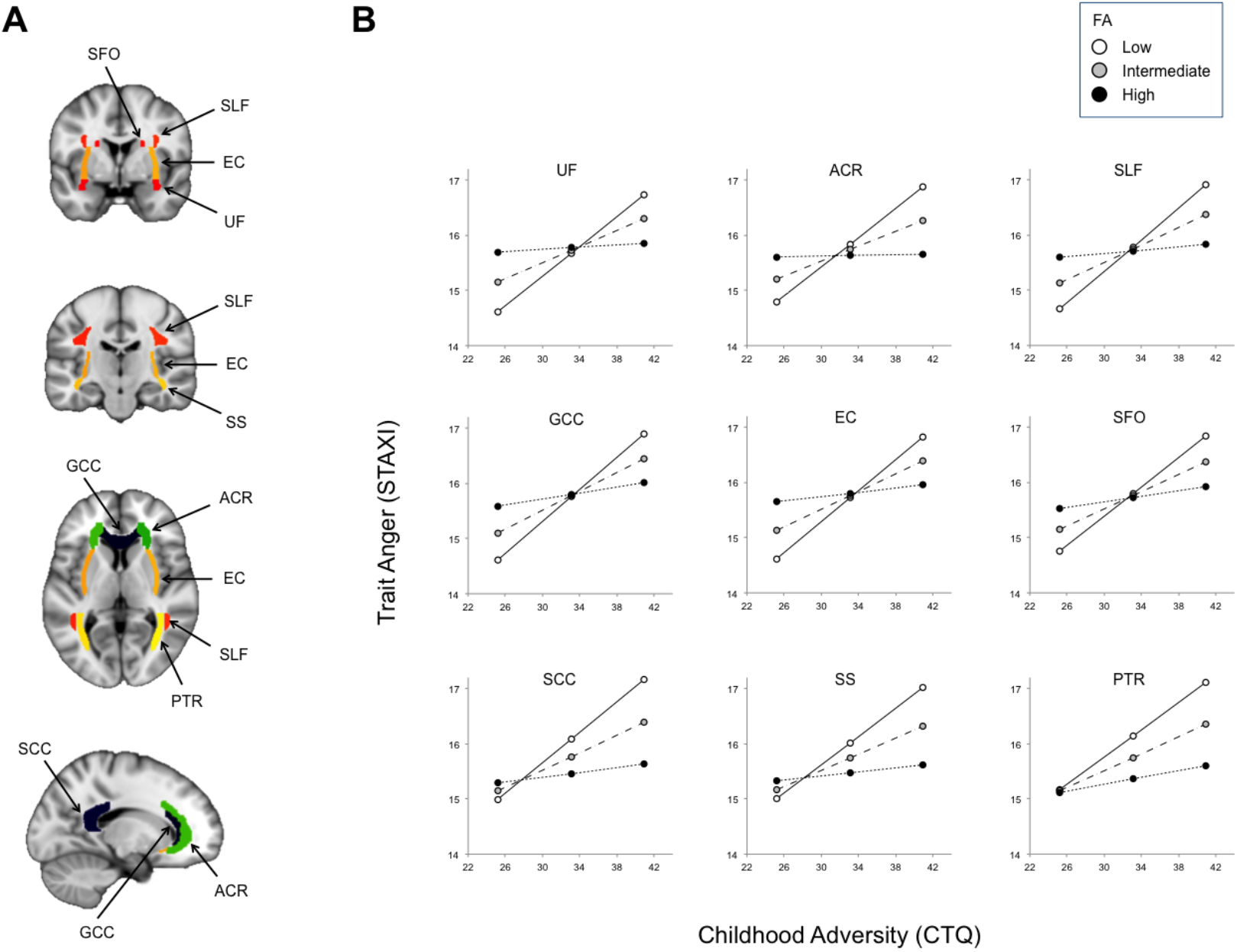
Widespread white matter pathway microstructural integrity as asssessed with dMRI moderates an association between childhood adversity and later trait anger. (A) The major fiber pathways exhibiting significant moderating effects (all *q*s < 0.05). (B) Positive association between childhood adversity and trait anger was attenuated in individuals with relatively higher microstructural integrity (i.e., FA) in these white matter pathways (black circles/dotted lines). PTR became significant when whole brain FA was included as a covariate. **Abbreviations**: FA (fractional anisotropy), ACR (anterior corona radiata), EC (external capsule), GCC (genu of the corpus callosum), PTR (posterior thalamic radiation), SCC (splenium of the corpus callosum), SFO (superior fronto-occipital fasciculus), SLF (superior longitudinal fasciculus), SS (sagittal stratum), UF (uncinate fasciculus).

## Discussion

Consistent with prior research and our primary hypothesis, we found that an association between childhood adversity and trait anger in young adulthood is moderated by the microstructural integrity of the UF, a corticolimbic pathway between the prefrontal cortex and amygdala also associated with the typical use of top-down cognitive reappraisal strategies to regulate negative emotion (Eden et al., 2015; Zuurbier et al., 2013). Surprisingly, however, 8 of the 24 white matter pathways examined yielded similarly significant effects, such that higher microstructural integrity attenuated the association between childhood adversity and trait anger. Notably, several of these other pathways including the anterior corona radiata (ACR), superior longitudinal fasciculus (SLF), and genu of the corpus callosum (GCC) also reflect prefrontal structural connections. These effects were relatively specific to trait anger, as none of the white matter pathways moderated the link between childhood adversity and trait anxiety; in other words, unlike trait anger, trait anxiety was strongly linked with childhood adversity regardless of the microstructural integrity of white matter pathways.

It is worth noting that the largest moderation effect was observed for the ACR, which is a major white matter pathway with reciprocal connections between the thalamus and the cerebrum that covers the prefrontal cortex and the anterior cingulate cortex, and has been suggested to be support multiple forms of information processing (Kelly et al., 2017; Wakana et al., 2004; Catani et al., 2002). Of particular relevance, studies have shown that individual differences in the microstructural integrity of the ACR is associated with executive control-related function such as (a) cognitive control, as measured by reaction times from a go/no-go task (Liston et al., 2006) or a modified Stroop task (Seghete et al., 2013); (b) executive attention, as measured by the time spent on resolving conflict during an attention network test (Niogi et al., 2010; Tang et al., 2010); and (c) working memory, as measured by the performance on the Memory for Digits subtest of the Wechsler Abbreviated Scale of Intelligence (Niogi and McCandliss, 2006). Furthermore, a recent large-scale meta-analysis across 4,322 individuals reported that the microstructural integrity of the ACR showed the largest deficits in schizophrenia, corroborating the findings from previous functional neuroimaging and behavioral studies demonstrating prefrontal executive function-related abnormalities in schizophrenia (Kelly et al., 2017).

The current finding with the ACR may be further relevant in the context of our previous functional neuroimaging study, where we observed that individuals with lower threat-related amygdala activity *and* higher executive control-related dlPFC activity had an attenuated association between childhood adversity and trait anger (Kim et al., in press). One possible interpretation of these converging findings is that higher microstructural integrity of prefrontal white matter enables dlPFC to exert more efficient control over other prefrontal and limbic areas, including the amygdala, thereby buffering against the negative effect of childhood adversity on trait anger. While we were unable to test this idea directly as our previous fMRI data set (Kim et al., in press) and current dMRI dataset have no overlap in participants, future investigations using concurrent structural and functional neuroimaging data can build on our findings.

In addition to the UF and ACR, which provide prefrontal structural connectivity, white matter pathways in other areas of the brain such as the external capsule (EC) and splenium of the corpus callosum (SCC) also showed significant moderating effects, implying that widespread white matter alterations may be involved in the emergence of trait anger associated with childhood adversity. However, it is important to consider the moderating effects of each pathway based on the structure of the interaction (Figure 1B). All interactions are based on the differences in slopes across varying FA levels (i.e., for all 8 pathways, association between childhood adversity and trait anger was attenuated in individuals with greater FA). Interestingly, some interactions are mostly driven by trait anger differences in individuals with higher childhood adversity, whereas others are additionally influenced by differences in individuals with lower childhood adversity. For the SS, SCC, and PTR, lower childhood adversity corresponded to lower trait anger regardless of the FA values in these pathways; as the level of childhood adversity increases, however, the relative integrity of these pathways becomes an important moderator. In this regard, these pathways fit the buffering account more so than others, as higher microstructural integrity appears to play a protective role against the association of childhood adversity and trait anger later in life. For the remaining pathways showing a significant moderating effect, the buffering account does not fully explain the observed interaction as individuals with higher microstructural integrity exhibited intermediate levels of trait anger, even when childhood adversity was low. Future studies utilizing a longitudinal design could further explore this unexpected effect, and therefore the present findings in these pathways should be interpreted with caution.

Our study, of course, is not without limitations that can be addressed in future research. First, as the data were collected via a cross-sectional design, causal inference is limited and interpretations of the findings should be accordingly tempered. Future work employing a longitudinal design would be able to mitigate the shortcomings associated with using retrospective reports to measure childhood adversity. Second, the data were acquired from a sample of high functioning university students, which limits the ability to generalize the findings to a broader population, especially to those with severe experiences of childhood adversity such as institutionalization, as well as those with pathological levels of anger or aggression. Those who experienced significant childhood adversity and still ended up as high functioning young adults, a general characteristic of individuals in a university study sample such as ours, may reflect their relative resilience to early life stress. Third, we note that as the fornix region is especially susceptible to partial volume effects (Pai et al., 2001; Prakash and Nowinski, 2006), the interpretation of the null finding within this pathway should be made with caution until replicated. We also note that the relatively small effect sizes reported here are consistent with previous neuroimaging studies of individual differences related to emotion (Zuurbier et al., 2013; Kim et al., 2017), likely reflecting the complex nature of variability that manifests in brain and behavior, which in turn highlights the importance of considering multiple moderating variables in future research. Finally, future studies could improve the quality of dMRI data by acquiring them with a higher angular sampling resolution (e.g., > 60 diffusion-weighted directions). That being said, we note that the FA data derived from a 15-direction vs. 61-direction dMRI dataset were comparable to one another in the context of their association with trait anxiety (Kim et al., 2017).

In summary, we report that an association between childhood adversity and trait anger later in life is moderated by individual differences the microstructural integrity of white matter pathways connecting multiple brain regions not limited to those within a corticolimbic circuit. Generally, higher white matter integrity was associated with an attenuated link between childhood adversity and trait anger. As such, the present findings provide further support for the importance of microstructural integrity of white matter pathways to mental health. If future investigations, especially those employing longitudinal designs, are able to replicate and extend these patterns to more diverse populations, such measures of white matter microstructural integrity may become a reliable and useful biomarker of an individual’s relative risk or resiliency to the experience of childhood adversity.

## Acknowledgements

This work was supported by Duke University and National Institutes of Health (NIH) Grants R01DA033369 and R01DA031579 to the Duke Neurogenetics Study; NIH Grant R01AG049789 to M.J.K., A.R.K., and A.R.H.; and the National Science Foundation Graduate Research Fellowship, Grant NSF DGE-1644868 to M.L.E. We thank the Duke Neurogenetics Study participants and the staff of the Laboratory of NeuroGenetics. In addition, the Brain Imaging and Analysis Center received support from the Office of the Director, National Institutes of Health under Award Number S10OD021480. The authors declare no competing financial interests.

